# Machine Learning-based Transcriptomic Data Analysis of Long COVID Patients: Generating Hypotheses for Molecular Signatures

**DOI:** 10.64898/2026.11.20.768015

**Authors:** John P. George, Shweta Rana, Dikscha Sapra, Harpreet Singh, Jyoti Sharma

**Affiliations:** Manipal Academy of Higher Education, Manipal, Karnataka 576104, India; Institute of Bioinformatics, International Technology Park, Bangalore 560066, India; Development Research Division, Indian Council of Medical Research, New Delhi 110029, India; Ram Lal Anand College, University of Delhi, New Delhi 110021, India

**Keywords:** post-COVID-19, Artificial intelligence, Transcriptome, Interaction Networks, Drugs

## Abstract

**Introduction:** Long COVID (LC) remains a global public-health challenge, affecting an estimated 10-30% of COVID-19-survivors. Machine learning (ML) can help identify molecular features associated with LC. Human ion channels (HICs), that regulate immune-cell signaling and cellular homeostasis, may contribute to LC-associated processes and offer insights into relevant pathways and potential drug-repurposing candidates.

**Methodology:** RNA-seq data from COVID-19-recovered and LC patients across six datasets were integrated, preprocessed, and corrected for batch effects. Supervised ML models were evaluated using leave-one-batch-out validation, with SHAP identifying features across validation folds. Functional enrichment, host-host and host-viral protein-protein interaction analyses were performed, followed by drug interaction analysis of potential HICs within the resulting interactome.

**Results:** A support vector classifier with an RBF kernel achieved a mean-AUC of 0.664 across six validation folds. SHAP analysis of the three selected folds (AUC = 0.722, 0.836, and 0.817) identified 225, 153, and 241 genes, respectively. Functional enrichment highlighted pathways including histidine metabolism and TNF signaling. Among the 613 genes identified across these folds, 9 were HICs and 39 were immune signaling-related genes (ISGs), whose protein-protein interaction analysis revealed a pharmacologically relevant interactome. Furthermore, four HICs including CLCN3, KCNQ3, RYR1, and ZACN interacted with 30 approved drugs.

**Conclusion:** ML-based analysis of integrated RNA-seq data identified HICs and ISGs associated with LC, while functional enrichment and PPI analyses provided insights into their potential biological relevance. Drug interaction analysis further highlighted candidate HIC-targeting drugs for future investigation in LC management. This computational framework provides preliminary, hypothesis-generating insights to guide future research.

## 1 Introduction

Nearly half a decade after the COVID-19 pandemic, long COVID (LC) has emerged as a major and heterogeneous public health concern. LC encompasses a broad spectrum of persistent or newly developed symptoms following acute SARS-CoV-2 infection, including fatigue, cognitive dysfunction, sleep disturbances, dyspnea, cough, musculoskeletal pain, and altered smell and taste [1], [2]. The current clinical and biological heterogeneity of LC suggests that no single pathological mechanism adequately explains its diverse manifestations. Proposed mechanisms include persistent immune dysregulation, altered host metabolism, endothelial dysfunction, mucosal barrier abnormalities, dysregulated erythropoiesis, hormonal disturbances, mitochondrial dysfunction, and, in some individuals, persistence of viral components [3], [4], [5]. These processes may interact with host-related factors such as age, sex, genetic background, comorbidities, vaccination status, and the timing and severity of the initial infection, further contributing to inter-individual heterogeneity [4].

Increasing evidence indicates that immune abnormalities can persist after the resolution of acute SARS-CoV-2 infection and are associated with LC [6], [7]. However, the immune alterations reported in LC are distinct from the acute inflammatory response’s characteristic of active SARS-CoV-2 infection. While acute infection may involve robust antiviral and inflammatory responses, LC has been associated with persistent alterations in multiple immune-cell populations and signaling pathways, although these changes vary across individuals and clinical phenotypes [4], [8]. Reported abnormalities in LC include T-cell exhaustion, altered regulatory T-cell (Treg) populations, expansion and activation of monocytes and neutrophils, increased levels of pro-inflammatory cytokines and chemokines, and alterations in levels of interleukin-18 (IL-18), interleukin-17 (IL-17) and further changes in naïve and effector T-cell, mucosal-associated invariant T (MAIT), and γδ T-cell populations [9], [10]. Impaired intestinal barrier integrity and microbial translocation have been associated with increased soluble CD14 (sCD14), while emerging evidence suggests that sCD14-associated, gamma-aminobutyric acid (GABA)- related signaling may promote T-helper 17 (Th17) differentiation and IL-17A production [11], [12]. Tumor necrosis factor (TNF)-related inflammation may also contribute to LC-associated immune alterations. Plasma-proteomic analyses have implicated TNF-α-associated signaling in vascular inflammation in LC [13], while a recent meta-analysis reported elevated circulating TNF-α in patients with post-acute sequelae of COVID-19, including beyond six months after infection [14]. These observations support the presence of sustained immune perturbation in a subset of individuals with LC, although the specific biological processes responsible for maintaining these alterations remain incompletely understood.

Several mechanisms, including viral persistence, immune dysregulation, altered erythropoiesis and red-blood-cell homeostasis, and endothelial dysfunction, have been proposed to explain the persistence and multisystem manifestations of LC, although their relative contributions remain incompletely understood [2], [15]. Viral RNA fragments or antigenic components have been detected in some individuals following acute infection; however, evidence remains insufficient to establish persistent, replication-competent SARS-CoV-2 as a universal mechanism of LC [16], [17], [18]. Viral persistence may therefore contribute to disease in specific patient subsets rather than represent a common explanation for LC pathogenesis. Emerging evidence also implicates altered erythropoiesis and hormonal imbalances in LC. Multiomics analyses have identified elevated heme-metabolism signatures and reduced androgenic steroids in individuals with LC, consistent with persistent inflammation, anemia, and stress erythropoiesis [19]. Furthermore, a study of individuals with LC and comorbid myalgic encephalomyelitis/chronic fatigue syndrome reported increased CD71-positive erythroid cells, disrupted erythropoiesis, including imbalanced sex hormones in males and females [20]. These findings support a potential link between inflammatory dysregulation, altered erythropoiesis, and endocrine disturbances, although their contributions to LC symptoms require further investigation [21]. Endothelial dysfunction, metabolic disturbances, and mucosal barrier disruption may also interact with immune dysfunction to influence disease manifestations [22], [23], [24]. Collectively, these findings support a multifactorial and heterogeneous model of LC in which different biological processes may coexist and vary across individuals.

Despite increasing evidence of immune and systemic abnormalities in LC, the molecular networks associated with these persistent changes remain incompletely characterized. In particular, the molecular interfaces connecting altered immune signaling with cellular homeostasis and host-pathogen interactions require further investigation. Transcriptomic profiling provides an opportunity to examine these molecular changes at a systems level and to identify patterns that may distinguish individuals with LC from individuals who have recovered from SARS-CoV-2 infection. Such analyses may help prioritize molecular features and biological pathways for subsequent experimental investigation.

Human ion channels (HICs) represent one potentially informative class of molecular regulators in this context. In addition to controlling membrane excitability and ion homeostasis, ion channels participate in intracellular signaling, immune-cell activation, cell proliferation, and other processes relevant to cellular responses to infection and inflammation [25], [26]. We previously identified associations between HICs, lipid metabolism, and SARS-CoV-2 infection and identified several HICs as potential candidates for pharmacological investigation [27]. We also investigated host responses associated with SARS-CoV-2–HIC interactions using machine-learning approaches and identified alterations in multiple host pathways [28]. These observations provided a rationale for examining ion channels in the context of LC, particularly in combination with immune signaling-related genes, while recognizing that their identification in computational analyses does not establish a causal role in disease.

Machine-learning and explainable artificial intelligence (AI) approaches provide complementary strategies for identifying transcriptomic features associated with complex clinical phenotypes. Recent studies have demonstrated the potential of AI-based approaches for identifying clinical and molecular patterns associated with LC [29]. However, models trained on heterogeneous transcriptomic datasets are susceptible to cohort-specific and technical variation and therefore require careful validation and cautious biological interpretation. Explainable AI methods such as SHapley Additive exPlanations (SHAP) can identify features that contribute to model predictions; However, the explained features represent model-associated signals rather than experimentally validated causal drivers or mechanistic mediators [30].

Here, integrated transcriptomic data analysis from multiple independent cohorts of COVID-19-recovered individuals and patients with LC were performed. Further, batch correction and leave-one-batch-out validation to evaluate the generalizability of supervised machine-learning models were applied. Subsequently explainable AI to identify genes contributing to classification across individual validation folds, with particular emphasis on HICs and immune signaling-related genes (ISGs) was applied. Functional enrichment analysis was performed to characterize the biological terms and pathways associated with the identified features, and drug interaction analysis was used to identify approved pharmacological interactions that may provide hypotheses for future investigation. These computational analyses are intended to generate testable hypotheses rather than establish causality, pathogenic mechanisms, or therapeutic efficacy.

Most likely, LC may be associated with a transcriptomic pattern involving HICs and ISGs that differ from the molecular state observed following recovery from SARS-CoV-2 infection. Therefore, the explainable machine learning-based analysis could prioritize reproducibly model-associated molecular features and biological pathways that provide candidate hypotheses regarding the immune and cellular alterations observed in LC.

## 2 Methods

### 2.1. Data acquisition and preprocessing

A list of 493 HICs was obtained from the Human Genome Organization Gene Nomenclature Committee (HGNC) database. Similarly, a list of 1,376 immune signaling-related genes (ISGs) was obtained from the InnateDB database on 05 November 2025 [31]. RNA-seq data comprising 96 COVID-19-recovered individuals and 122 LC patients were obtained from the National Center for Biotechnology Information (NCBI) Gene Expression Omnibus (GEO) and BioProject using the following accessions: GSE222253, GSE224615, GSE226260, GSE251849, GSE297499, and PRJNA1184095. The datasets were downloaded using the NCBI SRA Toolkit (version 3.0.6). Read quality was assessed using FastQC (version 0.11.9), followed by quality filtering and adapter trimming using Trimmomatic (version 0.39). The filtered reads were aligned to the human reference genome GRCh38.p13 using STAR aligner (version 2.7.11b), and the resulting alignment files were used for gene-level expression quantification with HTSeq (version 2.0.5). The overall methodology has been summarized in Figure 1.

**FIGURE 1.**
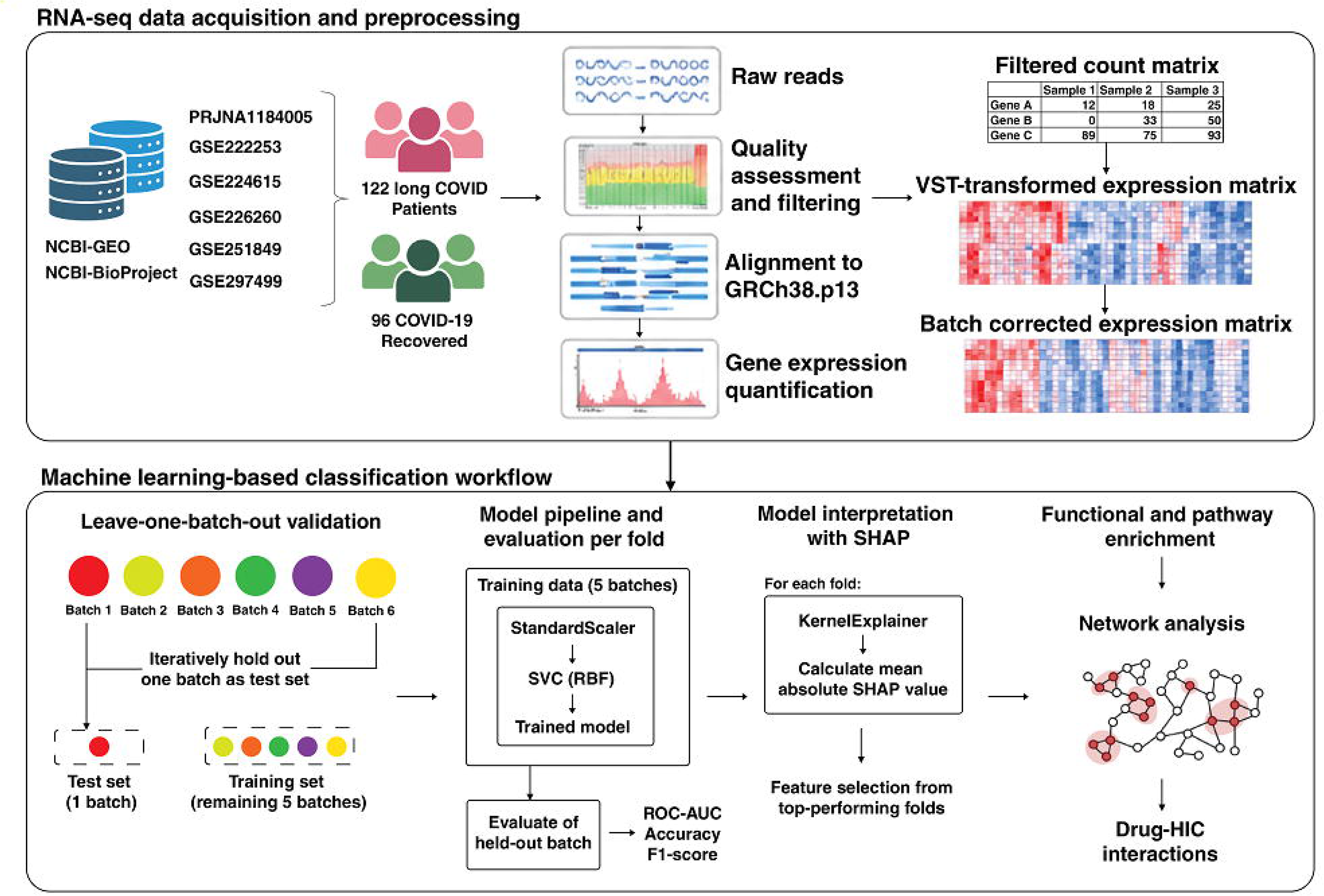
Workflow for machine learning-based transcriptomic data analysis of COVID-19 recovered and long COVID patients. The top panel illustrates data acquisition and preprocessing of the RNA-seq data while the bottom panel illustrates machine learning-based workflow consisting of leave-one-batch-out validation, feature extraction and their functional enrichment and biological relevance

As the datasets were generated by independent studies with differences in clinical characteristics, sampling time points, and sequencing protocols, inter-cohort heterogeneity was anticipated. Therefore, a uniform preprocessing workflow was applied to minimize technical variation and improve the consistency of downstream machine-learning analyses. The filtered raw count matrix was imported into DESeq2 (version 1.46.0) using the “DESeqDataSetFromMatrix” function. Variance-stabilizing transformation (VST) was subsequently performed using the “vst” function, which incorporates size-factor normalization based on the median-of-ratios approach to account for differences in sequencing depth and library size while reducing the dependence of variance on mean expression. The resulting transformed expression values were further subjected to batch-effect correction using the “removeBatchEffect” function from the limma package (version 3.62.2), retaining the biological condition in the model to minimize cohort-associated technical variation while preserving condition-related differences. The resulting expression matrix was used for downstream machine learning-based analyses.

### 2.2. Machine learning-based method workflow

A supervised machine learning-based workflow was implemented using scikit-learn (version 1.7.2) to distinguish COVID-19-recovered individuals from patients with LC. To evaluate model generalizability across independent cohorts and minimize the potential influence of cohort-specific characteristics, leave-one-batch-out (LOBO) cross-validation was employed. In each iteration, all samples from one batch (cohort) were held out as the test set, while samples from the remaining batches were used for model training. Thus, each batch served once as an independent held-out test set, and no samples from the held-out batch were used during model fitting or feature scaling.

Feature scaling was performed within the machine learning-based pipeline using StandardScaler(). The scaling parameters were estimated exclusively from the training samples in each LOBO iteration and subsequently applied to the corresponding held-out batch, thereby preventing information leakage during model evaluation. Model performance was evaluated on the held-out batch using receiver operating characteristic area under the curve (ROC-AUC), accuracy, balanced accuracy, sensitivity, specificity, F1-score, and confusion matrix.

For binary classification, true positive (TP) represented LC samples correctly classified as LC, whereas false positive (FP) represented COVID-19-recovered samples incorrectly classified as LC. True negative (TN) represented COVID-19-recovered samples correctly classified as recovered, and false negative (FN) represented LC samples incorrectly classified as recovered. Accuracy was calculated as the proportion of correctly classified samples among all evaluated samples:

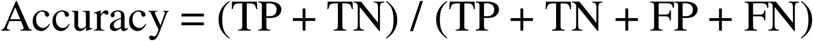

Sensitivity was calculated as the proportion of LC samples correctly identified by the model:

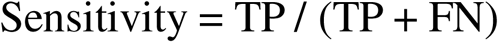

Specificity was calculated as the proportion of COVID-19-recovered samples correctly identified by the model:

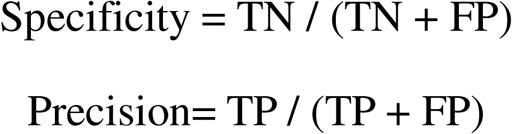

The F1-score was calculated as the harmonic mean of precision and sensitivity, and balanced accuracy was calculated as the mean of sensitivity and specificity. ROC-AUC was calculated using the model decision scores.

To interpret the contribution of individual genes to model predictions, SHapley Additive exPlanations (SHAP) were applied using KernelExplainer to each fitted LOBO model. A background set sampled exclusively from the corresponding training data was used for each fold. Thus, each held-out batch was associated with a fold-specific set of gene-level SHAP values. The magnitude of the SHAP value represented the strength of a gene’s contribution to the model output for an individual sample. To quantify the overall importance of each gene within a held-out batch, the mean absolute SHAP value across all samples in that fold was calculated and used to rank genes according to the magnitude of their contribution.

List of genes from top-performing LOBO folds that represented better classification of conditions were extracted for downstream interpretation. Further, SHAP-ranked genes were intersected with the predefined HICs and ISGs.

### 2.3. Functional enrichment analysis

Functional enrichment analysis for the identified features was performed using EnrichR. Gene Ontology (GO) terms were annotated for biological processes, molecular functions, and cellular components. Further, KEGG pathways enriched through the identified features were also identified.

### 2.4 Protein-protein interaction analysis

To identify the interactions between the identified HICs and immune signaling-related features, protein-protein interaction networks (PPINs) were constructed using STRING database [32]. Discarding the disconnected nodes, the interactions with a minimum required interaction score of 0.150 and false discovery rate of 5% extracted. Further, their interactions with SARS-CoV-2 proteins were parsed from BioGRID database [33]. The interactions between HICs, and immune signaling-related proteins with SARS-CoV-2 proteins were further visualized using Cytoscape (version 3.10.3) [34].

### 2.5. Drug interactions analysis

The identified HICs and ISGs were evaluated for their potential as therapeutic targets using the annotations from Drug-Gene Interaction Database (DGIdb) [35]. This led to identification of candidate genes with existing pharmacological modulators, thereby highlighting opportunities for drug repurposing and targeted therapeutic development for better management of disease.

## 3 Results

### 3.1. Model performance in leave-one-batch-out validation

The SVC with a radial basis function (RBF) kernel (C = 1.15; class weights balanced) was evaluated using a leave-one-batch-out validation strategy across six independent transcriptomic batches. Model performance varied across the held-out batches, with ROC-AUC values ranging from 0.452 to 0.836, accuracy from 53.8% to 87.5%, and balanced accuracy from 50.0% to 85.5%. The highest performance was observed when batch 4 (GSE251849) was held out, with an ROC-AUC of 0.836, accuracy of 87.5%, and balanced accuracy of 85.5%. Batch 5 (GSE297499) showed the second-highest performance, with an ROC-AUC of 0.817, accuracy of 76.9%, and balanced accuracy of 74.5%, followed by batch 1 (GSE222253), which achieved an ROC-AUC of 0.722, accuracy of 62.5%, and balanced accuracy of 62.5%. In contrast, batches 2, 3, and 6 showed limited discrimination, with ROC-AUC values of 0.579, 0.580, and 0.452, respectively (Table 1).

**Table 1:**
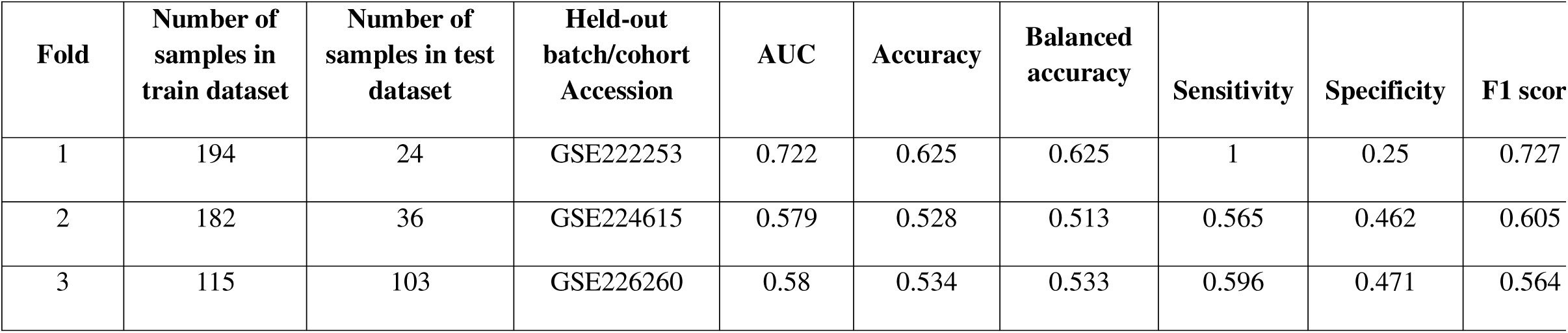
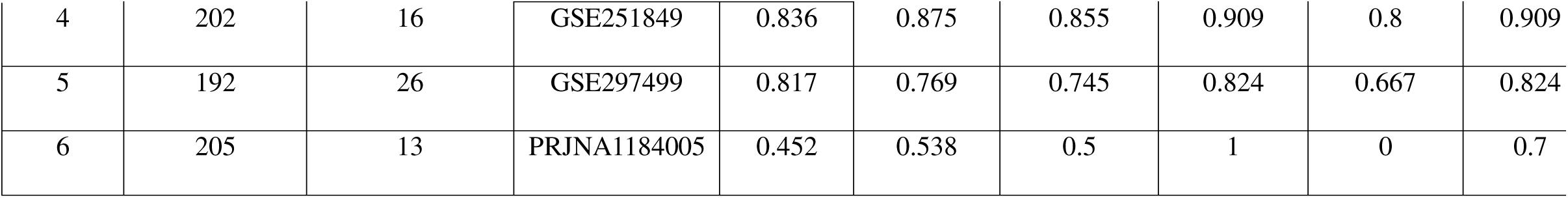
Leave-one-batch-out validation results for support vector classifier with RBF kernel.

| Fold | Number of samples in train dataset | Number of samples in test dataset | Held-out batch/cohort Accession | AUC | Accuracy | Balanced accuracy | Sensitivity | Specificity | F1 scor[illegible] |
|---|---|---|---|---|---|---|---|---|---|
| 1 | 194 | 24 | GSE222253 | 0.722 | 0.625 | 0.625 | 1 | 0.25 | 0.727 |
| 2 | 182 | 36 | GSE224615 | 0.579 | 0.528 | 0.513 | 0.565 | 0.462 | 0.605 |
| 3 | 115 | 103 | GSE226260 | 0.58 | 0.534 | 0.533 | 0.596 | 0.471 | 0.564 |
| 4 | 202 | 16 | GSE251849 | 0.836 | 0.875 | 0.855 | 0.909 | 0.8 | 0.909 |
| 5 | 192 | 26 | GSE297499 | 0.817 | 0.769 | 0.745 | 0.824 | 0.667 | 0.824 |
| 6 | 205 | 13 | PRJNA1184005 | 0.452 | 0.538 | 0.5 | 1 | 0 | 0.7 |

Across all six validation folds, the SVC-RBF model achieved a mean ROC-AUC of 0.664 ± 0.152, mean accuracy of 64.5% ± 14.6%, and mean balanced accuracy of 62.9% ± 14.4%. The mean sensitivity was 81.6% ± 19.4%, whereas the mean specificity was 44.1% ± 28.7%, indicating greater classification of LC samples than recovered samples across the validation folds. The mean F1 score was 72.1% ± 13.0%. ROC curves and confusion matrices for top-performing folds are summarized in Figure 2.

**FIGURE 2.**
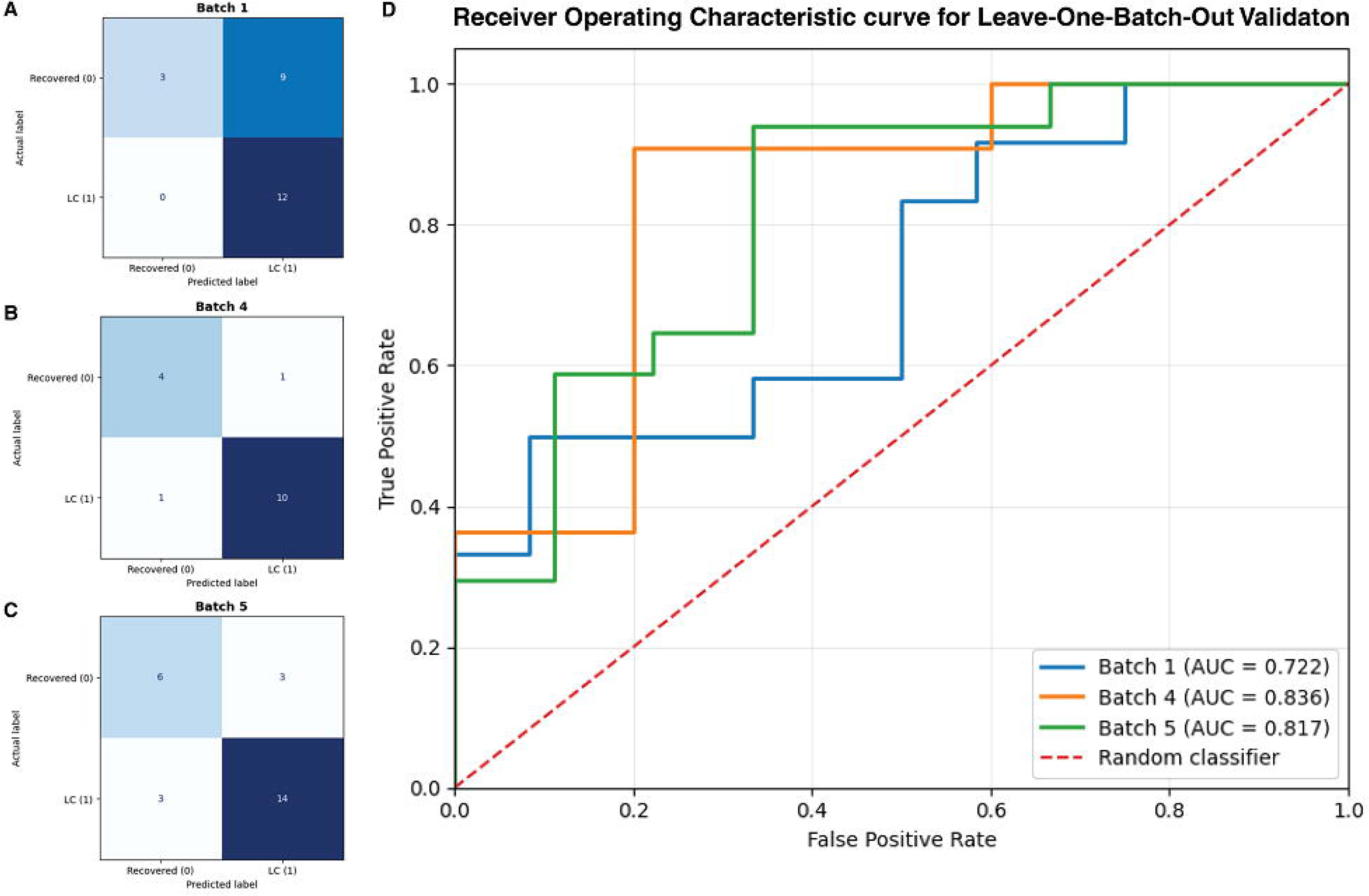
Confusion matrices for the top-performing validation folds (A: batch 1, B: batch 4, and C: batch 5) and (D) receiver operating characteristic curves for the three selected batches

### 3.2. Feature explanation and biological interpretation using Explainable Artificial Intelligence (AI)

Explainable AI was subsequently applied to the SVC-RBF model to characterize the transcriptomic features contributing to classification in the leave-one-batch-out validation analysis. SHAP values were calculated separately for the three validation folds that showed the highest ROC-AUC values (batches 4, 5, and 1) (Supplementary File 1). Features identified across these folds were combined to generate a non-redundant feature set for downstream biological interpretation.

A total of 613 unique genes were identified from the three selected validation folds. Among these, 9 genes were classified as HICs - *CLCN3, CNGA4, CRACR2B, KCNQ3, KCTD17, RYR1, SLC16A13, SLC8A3, ZACN*, while 39 were identified as ISGs - *ABL1, ACTR3, AKIRIN2, AKNA, ANGPT1, ANKRD17, C4BPB, C8G, CALCOCO2, CD19, CD22, DNM1L, FGR, FOS, IL18R1, IRGM, ITCH, LAT2, LY96, MDM2, NR1H3, NR3C1, OAS2, P2RY14, PRKCE, RICTOR, RPS6KA5, RUSC1, SERPINB9, SIGLEC14, SIGLEC6, SIKE1, SLC15A4, TLR8, TPT1, TXN, UBA52, USP2, VDR* (Figure 3). Mean absolute SHAP values were used to quantify the overall contribution of individual features to the model predictions, with higher values indicating a greater contribution to the classification output. The identified features therefore represent genes that were associated with the model’s ability to distinguish LC from COVID-19-recovered samples within the selected validation folds (Supplementary File 1).

**FIGURE 3.**
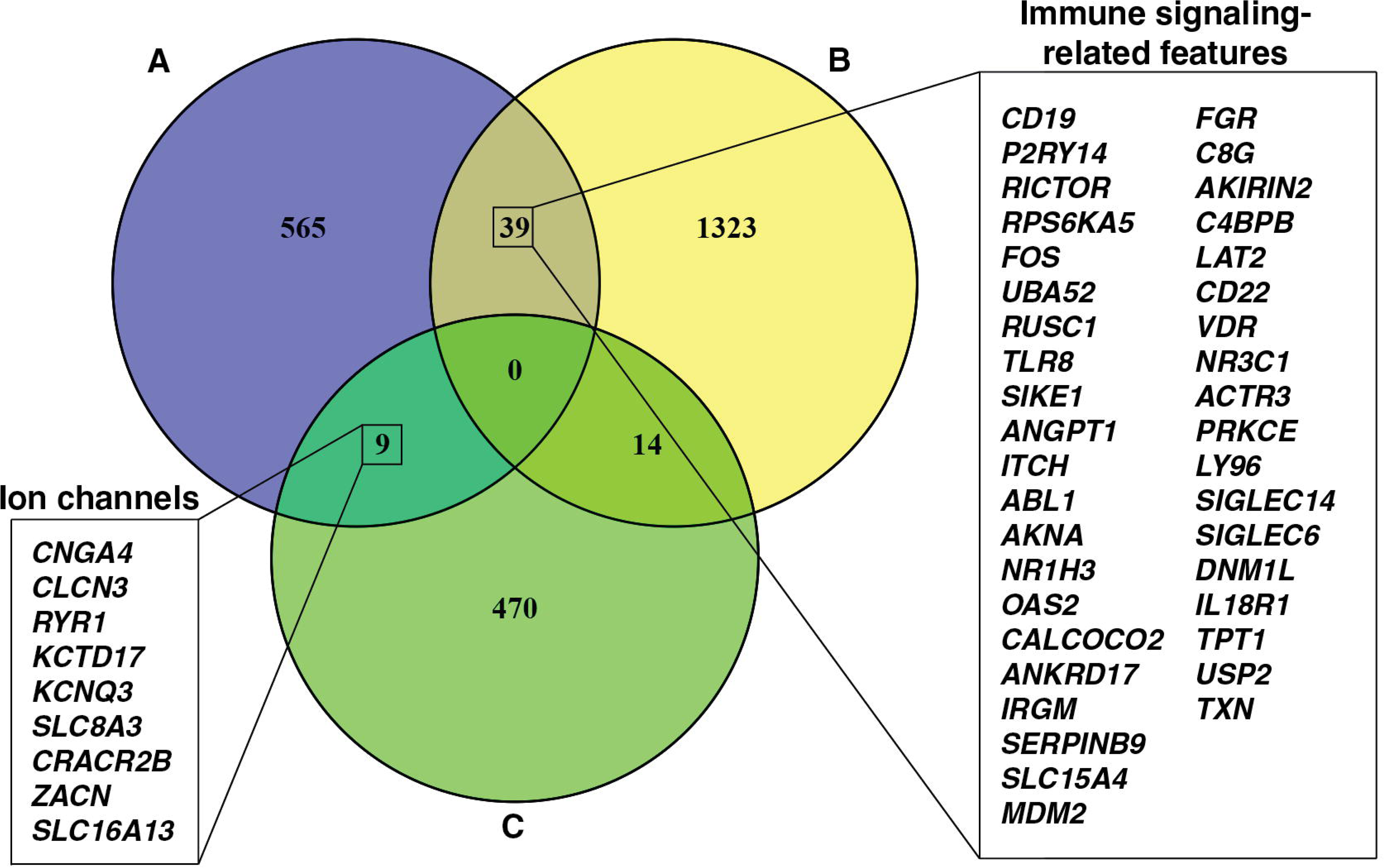
Overlap between (A) features from top performing batches (1, 4, and 5), (B) immune signaling-related features, and (C) human ion channels (HICs)

### 3.3 Functional and pathway enrichment analysis

Functional enrichment analysis of the 613 non-redundant genes identified significantly enriched biological processes, molecular functions, and cellular components. Among the biological processes, several immune-related processes were enriched, including regulation of cytoplasmic pattern-recognition receptor signaling, Fc-gamma receptor signaling involved in phagocytosis, Fc receptor-mediated stimulatory signaling, regulation of B-cell receptor signaling, and regulation of cell activation. Processes related to hematopoietic regulation, including regulation of hematopoietic stem cell differentiation and regulation of hematopoietic progenitor cell differentiation, were also significantly enriched, along with iron ion transport. Mitochondrial and cellular stress-related processes, including positive regulation of mitophagy, regulation of mitophagy, positive regulation of mitochondrial fission, regulation of mitochondrial membrane permeability, positive regulation of autophagy of mitochondrion, and regulation of cellular response to oxidative stress, were also significantly enriched. In addition, neurotransmitter-related processes, including cellular response to catecholamine stimulus, response to dopamine, cellular response to dopamine, and regulation of neurotransmitter uptake, were identified (Figure 4A).

**FIGURE 4.**
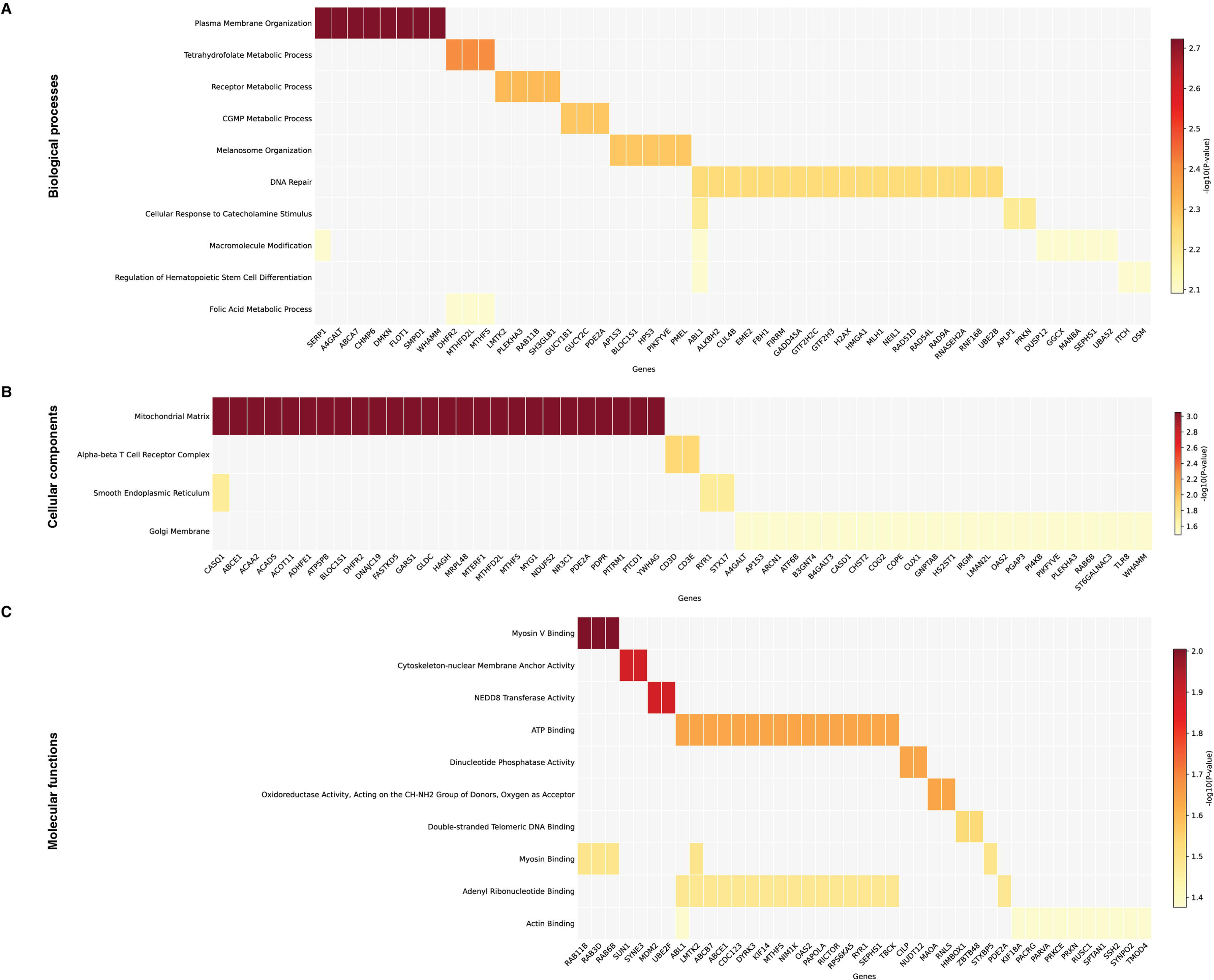
Functional enrichment analysis of 613 features identified in the top performing batches (1, 4, and 5). A, B, and C correspond to biological processes, cellular components, and molecular functions, respectively

For cellular components, significant enrichment was observed for the mitochondrial matrix, alpha-beta T-cell receptor complex, smooth endoplasmic reticulum, and Golgi membrane (Figure 4B). For molecular functions, significant enrichment was observed for actin binding, myosin binding, cGMP binding, ATP binding, and adenyl ribonucleotide binding, representing cytoskeletal, cyclic-nucleotide-associated, and nucleotide-binding functions (Figure 4C).

Pathway enrichment analysis further identified three significantly enriched pathways including histidine metabolism, arginine and proline metabolism, and the TNF signaling pathway. Collectively, the enrichment analyses indicated that the features were distributed across biological processes and pathways related to immune signaling, hematopoietic regulation, mitochondrial and metabolic processes, neurotransmitter-related functions, cytoskeletal activity, and membrane-associated cellular compartments. A pathway map was generated to highlight the identified features in TNF signaling and histidine metabolism pathway (Figure 5)

**FIGURE 5.**
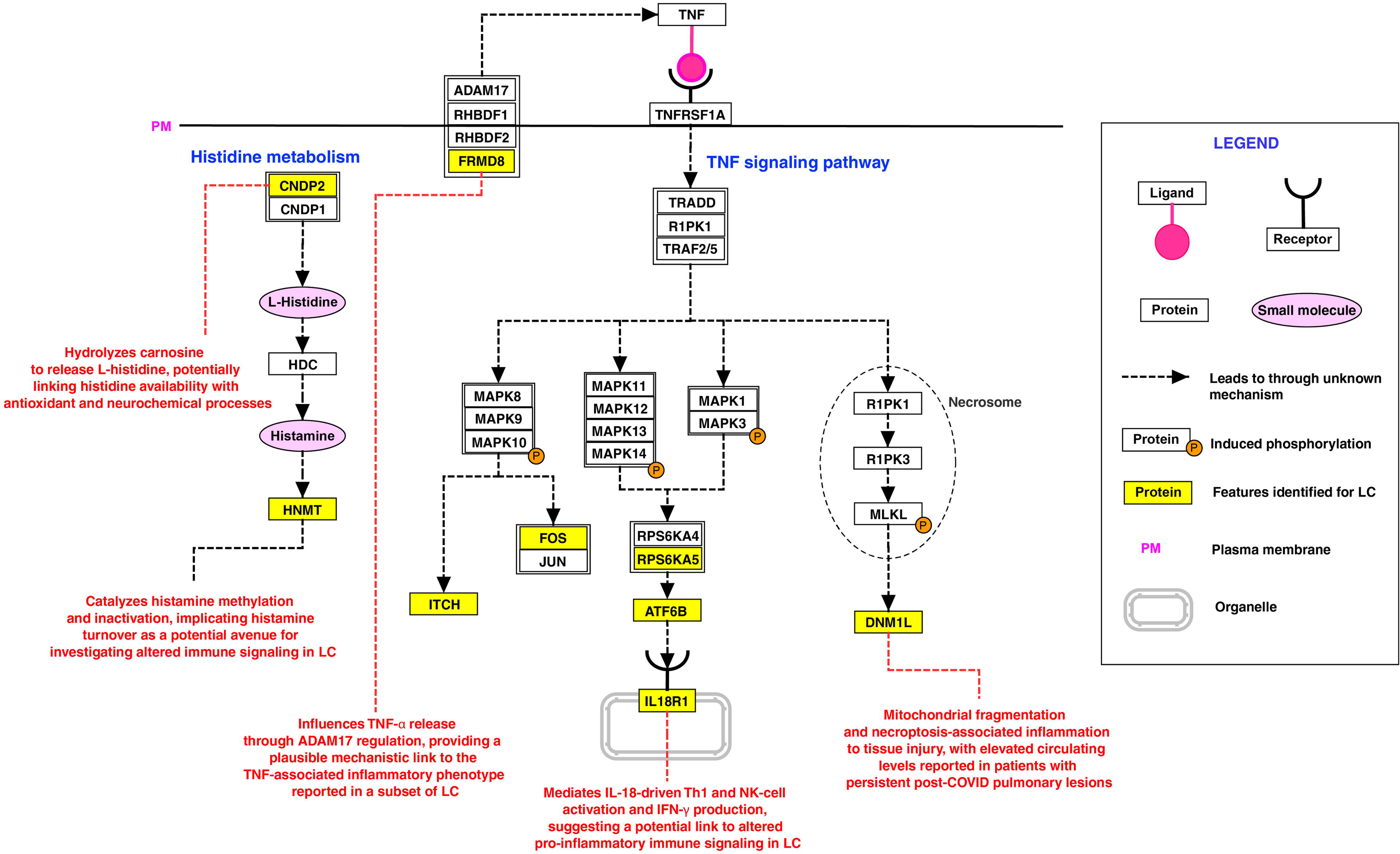
Pathway map consisting of TNF signaling and histidine metabolism pathway. The nodes highlighted yellow are the identified features through machine learning-based approach

### 3.4. PPIN between HICs, ISGs and SARS-CoV-2 proteins

The PPI network derived from the STRING database revealed predicted PPIs involving 7 HICs and 37 immune signaling-related proteins. Among these, 21 host proteins, comprising 18 immune signaling-related proteins (ACTR3, AKNA, ANKRD17, C4BPB, CALCOCO2, DNM1L, IL18R1, ITCH, MDM2, NR3C1, RICTOR, SERPINB9, SIKE1, SLC15A4, TPT1, TXN, UBA52, and USP2) and 3 HICs (CLCN3, KCNQ3, and RYR1), were predicted to interact with 27 SARS-CoV-2 proteins, including S, E, M, N, ORF10, ORF14, ORF3a, ORF3b, ORF6, ORF7a, ORF7b, ORF8, ORF9b, ORF9c, NSP1, NSP11, NSP12, NSP14, NSP15, NSP16, NSP2, NSP3, NSP4, NSP5, NSP6, NSP8, and NSP9 (Figure 6).

**FIGURE 6.**
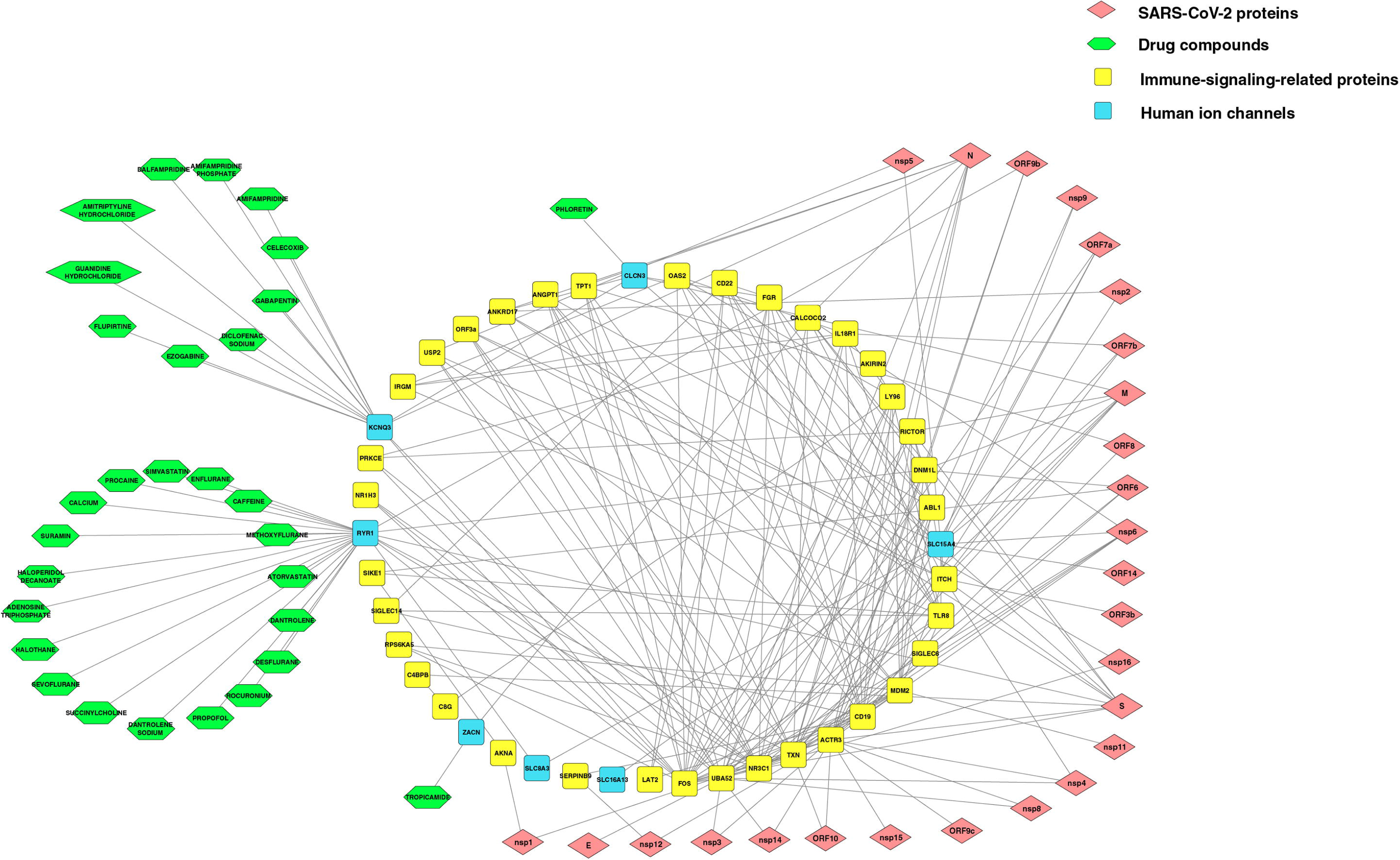
Protein-protein interaction between human ion channels (Blue), immune signaling-related proteins (Yellow), and further with SARS-CoV-2 proteins (Red). Drugs interacting with ion channels are represented in green color hexagonal node

### 3.5. Drug-gene interactions

Among the seven HICs identified in the PPIN, five HICs (CLCN3, KCNQ3, RYR1, SLC8A3, and ZACN) were associated with interactions involving 58 drug compounds, resulting in a total of 62 drug-gene interactions. Of these interactions, CLCN3, KCNQ3, RYR1, and ZACN were found to interact with 30 approved drugs (Figure 6).

## 4 Discussion

LC is a heterogeneous post-infectious condition in which persistent immune, metabolic, vascular, neurological, and cellular abnormalities may coexist across patients. Here, we integrated RNA-seq data from six independent cohorts of COVID-19-recovered individuals and patients with LC and evaluated an SVC-RBF model using leave-one-batch-out validation. Rather than observing uniformly high classification performance, we found substantial variation across held-out batches, with the highest ROC-AUC values observed for batches 1, 4, and 5 and limited discrimination in batches 2, 3, and 6. This variability is important in the context of LC heterogeneity and indicates that transcriptomic differences between recovered individuals and LC patients are not uniformly represented across independent cohorts. Previous transcriptomic studies have similarly demonstrated molecular differences between LC and recovered individuals and have identified heterogeneous immune and metabolic signatures among patients with LC [36], [37].

The mean ROC-AUC of 0.664 and balanced accuracy of 0.629 observed across all six batches suggest that the model captured a detectable but moderate transcriptomic signal rather than a universally discriminative LC signature. In particular, the stronger performance in batches 1, 4, and 5 provided a basis for subsequent feature interpretation, whereas the near-chance performance of batches 2, 3, and 6 demonstrates that the identified signal may be influenced by cohort-specific biological or technical characteristics. Batch 6 further illustrated this limitation, as all samples were classified as LC, resulting in a sensitivity of 1.00 but specificity of 0.00. Thus, the present model should not be interpreted as a diagnostic classifier for LC. Instead, the leave-one-batch-out analysis provides evidence that some transcriptomic patterns can generalize across independent cohorts while also highlighting the substantial inter-cohort variability inherent to LC datasets. Differences in gender, clinical phenotype, sampling time, disease severity, vaccination history, viral exposure, and other cohort characteristics have previously been associated with variation in LC-associated immune signatures [38], [37].

SHAP analysis of the three higher-performing validation folds identified 613 non-redundant genes, including 9 HICs and 39 ISGs. The use of mean absolute SHAP values allowed the magnitude of each feature’s contribution to model prediction to be quantified across samples. Importantly, previous studies have also used machine learning and transcriptomic approaches to identify candidate molecular signatures associated with LC, including studies based on recovered-versus-LC comparisons and broader multi-dataset analyses [36], [39]. The present analysis differs in its emphasis on a predefined set of HICs and ISGs, combined with cohort-aware validation and fold-specific SHAP interpretation. This framework provides a means of prioritizing genes for downstream biological analyses while retaining the exploratory nature of the study.

The identification of HICs among the model-associated features is of particular interest. Ion channels regulate ion homeostasis, membrane potential, intracellular signaling, and cellular activation and can therefore influence processes occurring across immune, metabolic, vascular, and neuronal systems. Although the specific channels identified in the present study require experimental validation, independent studies provide evidence that ion channel dysfunction can occur in post-COVID conditions [27], [40]. For example, altered TRPM3-dependent calcium signaling has been reported in NK cells from individuals with post-COVID condition, and subsequent work has examined restoration of TRPM3 channel function as a potential therapeutic avenue [41], [42], [43].

The functional enrichment results provided additional biological context for the model-associated genes. Immune-related processes, including pattern-recognition receptor signaling, Fc-gamma receptor-mediated phagocytosis, Fc receptor signaling, B-cell receptor regulation, and cell activation, were prominently represented. These findings align with accumulating evidence that LC is associated with persistent remodeling of both innate and adaptive immunity, including altered monocyte phenotypes, T-cell exhaustion, and changes in other lymphocyte populations [44], [45]. The enrichment of hematopoietic stem- and progenitor-cell differentiation and iron transport is also notable considering reported abnormalities in erythropoiesis, heme metabolism, and inflammatory states associated with LC [44], [19]. In the Immunophenotyping Assessment in a COVID-19 Cohort, a multiomics recovery factor associated with LC was linked to inflammatory mediators, heme-metabolism signatures, reduced lymphocyte frequencies, and increased myeloid-cell frequencies, supporting the relevance of immune and hematopoietic alterations to persistent post-COVID states [19].

Mitochondrial and cellular-stress-related enrichment represented another prominent feature of the analysis. Mitophagy, mitochondrial fission, mitochondrial membrane permeability, mitochondrial matrix localization, and oxidative-stress-related processes were among the enriched terms. These findings are compatible with growing evidence of altered mitochondrial function, bioenergetics, and oxidative metabolism in LC, particularly in relation to fatigue and exercise intolerance [5], [46]. Additionally, enrichment of neurotransmitter- and catecholamine-related processes provides insights on neurological and autonomic manifestations in LC [47], [48]. These observations should nevertheless be regarded as functional associations, as bulk blood transcriptomic enrichment cannot establish whether the corresponding biological changes occur directly within affected neuronal tissues.

The enrichment of the TNF signaling pathway, together with histidine metabolism and arginine and proline metabolism, further indicates that inflammatory and metabolic processes are represented within the model-associated gene set. Persistent inflammatory protein signatures have previously been described in subsets of individuals with LC, including an inflammatory phenotype involving TNF-associated signaling [49]. Recent systems-level analyses have also identified metabolic disturbances involving amino-acid pathways in LC and related post-infectious conditions [50]. The concordance of these pathway-level findings with the functional enrichment analysis suggests that the identified genes are likely distributed across immune and metabolic processes.

A pathway map was generated to represent model-associated genes overlaid onto curated pathways, including TNF signaling and histidine metabolism (Figure 5). Within histidine metabolism, CNDP2 and HNMT link the identified features to dipeptide metabolism and histamine degradation, respectively. Histamine-related mechanisms have been proposed in LC, although supporting evidence remains mixed [51], [52], [53]. Within TNF signaling, FRMD8 stabilizes the iRhom-ADAM17 complex involved in TNF-α shedding [54], while FOS and RPS6KA5 are associated with MAPK-mediated signaling responses. Other highlighted features, including ITCH, ATF6B, IL18R1, and DNM1L, suggest potential links to signaling regulation, cellular stress, cytokine responses, and mitochondrial dynamics. These findings provide additional context for reported TNF-associated inflammation and metabolic dysregulation in LC [49], [50].

The PPI analysis further placed a subset of the identified HICs and immune signaling-related proteins within a broader host-virus interaction network. Seven HICs and 37 immune signaling-related proteins were represented in the STRING network, and 21 host proteins were connected to 27 SARS-CoV-2 proteins through interactions obtained from BioGRID. Furthermore, the host-SARS-CoV-2 interactions obtained from interaction databases may originate from studies of acute infection and other experimental systems. Evidence for persistent SARS-CoV-2 in LC remains limited and inconsistent, with some studies not detecting residual circulating virus months after acute infection [16], [37].

The drug-gene interaction analysis identified 58 compounds associated with five HICs, corresponding to 62 drug-gene interactions, of which four HICs interacted with 30 approved drugs. These findings demonstrate that some of the identified HICs have existing pharmacological annotations and therefore provide hypothesis for drug-repurposing in LC. However, database-level drug-gene interactions do not establish the efficacy of a drug in LC. Computational drug-repurposing strategies are most appropriately viewed as candidate-prioritization approaches that require subsequent pharmacological and clinical validation [55].

## Limitations

Several limitations should be considered when interpreting these findings. First, the study was based on bulk RNA-seq transcriptomic data obtained from blood-derived samples, which capture gene expression patterns but do not directly verify protein-level changes. Nevertheless, these findings provide an initial line of evidence for future studies aimed at establishing the relationship between transcriptomic changes, protein-level alterations, and LC. Second, the six datasets differed in clinical characteristics, sampling time points, sequencing procedures, sex distribution, vaccination history, viral exposure, and LC phenotype. This heterogeneity is reflected in the substantial variation in model performance across the held-out batches and limits the extent to which a universal LC transcriptomic signature can be inferred. Third, SHAP values quantify the contribution of features to model predictions but do not establish their causality in the disease. Fourth, pathway enrichment and network mapping analyses are inherently dependent on existing pathway annotations and gene-set databases. Enrichment reflects statistical over-representation rather than causal involvement, and pathway maps presented here should be regarded as hypothesis-generating frameworks. Functional validation using experimental perturbation models will be necessary to establish mechanistic relevance. Further, the present study does not establish the direction of gene dysregulation or provide sufficient evidence to predict the therapeutic directionality or efficacy of the identified drug interactions in LC. Further experimental and clinical studies are required to validate these interactions and elucidate their downstream biological effects before their therapeutic potential for LC management can be established.

## 5 Conclusion

Cohort-aware machine learning-based approach led to the identification of HICs and ISGs, together with their enrichment across immune, hematopoietic, mitochondrial, metabolic, neurobiological, and membrane-associated processes, provides testable hypotheses regarding molecular alterations associated with LC. The identified HICs and their pharmacological annotations provide a foundation for future functional studies to investigate potential associations between specific ion channel pathways and LC phenotypes. However, further experimental and clinical validation is necessary to establish the biological significance and therapeutic relevance of these findings in LC management.

## Acknowledgements

We would like to thank the authors of the manuscripts for making the datasets used in this study publicly available. J.S. would like to thank Council of Scientific and Industrial Research (CSIR), Government of India [37WS(0114)/2023-24/EMR-II/ASPIRE] and Indian Council of Medical Research (ICMR), Government of India [FIW-2025-01-00001091] for the research support J.P.G. is supported by the CSIR [37WS(0114)/2023-24/EMR-II/ASPIRE]. J.S. was a recipient of the Bio-CARe Women Scientist award from the Department of Biotechnology (DBT), Government of India.

## CRediT authorship contribution statement

John P. George: Writing – review & editing, Writing – original draft, Visualization, Methodology, Investigation, Formal analysis, Data curation. Shweta Rana: Writing – review & editing. Dikscha Sapra: Writing – review & editing. Harpreet Singh: Writing – review & editing, Supervision. Jyoti Sharma: Writing – review & editing, Supervision, Methodology, Investigation, Funding acquisition, Conceptualization.

## Data availability

The datasets used in this study are available in NCBI’s Gene Expression Omnibus database (https://www.ncbi.nlm.nih.gov/geo/) and BioProject (https://www.ncbi.nlm.nih.gov/bioproject/). These datasets were derived using the following accessions - GSE222253, GSE224615, GSE226260, GSE251849, GSE297499, and PRJNA1184095. The pathway map is available in GPML format through the GitHub repository via the following URL: https://github.com/js-iob/LongCovid_MachineLearning/blob/main/LC_PW_Map_101126.gpml. Customized R and Python source codes are available through the following URL: https://github.com/js-iob/LongCovid_MachineLearning on 16 October 2026.

---

**SUPPLEMENTARY FILE 1**- Model-associated features identified using SHAP across all six batches

